# Semi-*in vitro* Reconstitution of Roseocin, a Two-Component Lantibiotic from an *Actinomycete*

**DOI:** 10.1101/464644

**Authors:** Mangal Singh, Dipti Sareen

## Abstract

Lantibiotics are lanthionine containing peptide natural products that belong to the class of ribosomally synthesized and post-translationally modified peptides (RiPPs). Recent expansion in the availability of microbial genomic data and *in silico* analysis tools have accelerated the discovery of these promising alternatives to antibiotics. Following the genome-mining approach, a biosynthetic gene cluster for a putative two-component lantibiotic roseocin was identified in the genome of an *Actinomycete, Streptomyces roseosporus* NRRL 11379. Post-translationally modified lanthipeptides of this cluster were obtained by heterologous expression of the genes in *E. coli,* and were *in vitro* reconstituted to their bioactive form. The two lanthipeptides displayed antimicrobial activity against Gram-positive bacteria only synergistically, a property reminiscent of two-component lantibiotics. Structural analysis of the α-component identified a disulfide bridge flanking two of its four thioether bridges and the β-component having six thioether bridges with its C-terminus extended than the previously known two-component lantibiotics.

## INTRODUCTION

Many of the clinically used antibiotics are derived from the genus *Streptomyces.* It is the largest genus of *Actinobacteria*, Gram-positive and has a high GC content genome. The arsenal of antimicrobial compounds that this genus produces includes polyketides, non-ribosomal, and ribosomal peptides. Recent studies have focussed on the discovery of ribosomally synthesized and post-translationally modified peptides (RiPPs) in light of the need for novel antimicrobials to combat the rising antimicrobial drug resistance (Basi-Chipalu, 2016). A particular focus has been on the discovery of novel lantibiotics (a class of RiPPs) that display their activity against the drug-resistant bacteria. Here, we have heterologously produced and characterized a lantibiotic roseocin from *Streptomyces roseosporus* NRRL 11379, by employing a semi-*in vitro* reconstitution approach.

As with the polyketides and non-ribosomal peptide antibiotics, lantibiotics are also synthesized from a biosynthetic gene cluster (BGC). A genetically encoded lantibiotic is initially synthesized as a linear peptide (precursor peptide) with two important regions, an N-terminal leader peptide region separated by a proteolytic cleavage site from its C-terminal core-peptide region that is rich in Cys/Ser/Thr residues. While the leader peptide guides the precursor peptide to different enzymes, the core-peptide forms the final lantibiotic structure. Lantibiotics are synthesized in two steps by (1) dehydration of Ser/Thr residues in the core-peptide followed by (2) intra-peptide Michael addition of cysteine residues to form lanthionine or thioether bridges (cyclization). These lanthionine rings are installed by various enzymes which define the four classes of lantibiotics. In class I lantibiotics, two separate enzymes for dehydration (LanB) and cyclization (LanC) are encoded in the BGC along with the gene for the precursor peptide (LanA). Besides this, a lantibiotic BGC encodes a protease (LanP) for the proteolytic removal of the leader peptide, a transporter (LanT) for extracellular transport of the modified peptide, immunity proteins (LanF, LanE, LanG and sometimes LanI) and a two-component response regulation system (LanR and LanK). Class II, III and IV lantibiotic BGCs encode a single lanthionine synthetase instead of separate dehydratase and cyclase enzymes for the lanthionine ring formation. The class III and IV lantibiotics have bioactivities majorly other than antimicrobial action like morphogenetic(Ueda *et al.*, 2002), antiviral, antiallodynic(Férir *et al.*, 2013), antinociceptive (Iorio *et al.*, 2014) and antidiabetic activities(Iftime *et al.*, 2015). The class II lantibiotics with a single lanthionine synthetase are antimicrobial and also the most heterologously characterized lantibiotics.

In class II lantibiotics, lanthionine ring installation in the core-peptide is done by a single lanthionine synthetase LanM, followed by the concomitant proteolytic removal of the leader and extracellular transport by a bifunctional transporter (LanT_p_). The promiscuity of LanM in lanthionine ring installation on its precursor peptides can be low (dedicated LanM for a single peptide, as in haloduracin) to very high (single LanM acting on >17 precursor peptides in prochlorosins). Class II lantibiotics are further of two types depending upon the constituent peptides and their bioactivity. Single component lantibiotics contain peptides which are all antimicrobial (Wang *et al.*, 2014) and the two-component lantibiotics contain two different types of peptides that show synergistic antimicrobial activity. Most of the two-component lantibiotics have a lipid II binding α-peptide along with a β-peptide, which acts in synergy with the α-peptide, thereby enhancing its activity manifolds (Bakhtiary *et al.*, 2017). Two-component lantibiotics are synthesized either by a single LanM which installs lanthionine rings on these two different types of peptide i.e. both on the α and β precursor peptide (Booth *et al.*, 1996; Lohans *et al.*, 2014; Huo and van der Donk, 2016), or by two separate LanM enzymes, each of which is specific to either the α or the β precursor peptide(s) (McClerren *et al.*, 2006; Caetano *et al.*, 2011; Zhao and van der Donk, 2016). While there are many examples of characterized and putative two-component lantibiotics (Singh and Sareen, 2014; Zhang *et al.*, 2015) synthesized by two LanMs, only three are known to be processed by a single LanM i.e. carnolysin and cytolysin which are homologs (Booth *et al.*, 1996; Lohans *et al.*, 2014) and bicereucin having D-amino acids as the major post-translational modification compared to lanthionine (Huo and van der Donk, 2016).

Understanding of genetic organization of lantibiotic encoding BGCs has led to the discovery of novel lantibiotics by combinatorial approach of genome mining, activity-based screening of the thus identified potential producers and/or *in vitro* reconstitution of BGCs from the native producers (McClerren *et al.*, 2006; Begley *et al.*, 2009). Till date, two-component lantibiotics have been isolated and are confined to only *Firmicutes* (Zhang *et al.*, 2015). In our earlier genome mining study (Singh and Sareen, 2014), we identified three putative two-component lantibiotic clusters in *Actinomycetes* having two precursor peptides and two LanMs, following a microbial genome database mining strategy for novel bifunctional transporter LanT_p_. In addition, a cluster identified in *S. roseosporus* NRRL 11379, encoded for a single LanM to process two precursor peptides. The presence of six cysteine residues in each of the two peptides of this cluster led us to speculate that they might form a highly constrained structure with difference in bioactivity than the existing lantibiotics. Additionally, we wanted to analyze whether these peptides show synergistic, as in two-component lantibiotics, or additive bioactivity. Hence, this cluster was undertaken for characterization by a semi-*in vitro* reconstitution approach, involving heterologous production of the lanthipeptides in *E. coli* (Shi *et al.*, 2011).

## RESULTS

### Bioinformatic analysis of Roseocin as a two-component lantibiotic

The roseocin cluster was identified following a genome mining study for novel HalT homologs (Singh and Sareen, 2014). The roseocin biosynthetic gene cluster comprises of putative genes for the two precursor peptides (RosAs), a lanthionine synthetase (RosM), a bifunctional transporter (RosT_p_), immunity proteins (RosEFG) and a transcriptional regulator (Figure 1 and Table S1). The two RosA peptides display an overall 68% similarity (40% identity) in the 75 amino acid overlap (Figure 1B). Although the leader peptide is highly similar, the core-peptide region is variable in the composition of lanthionine forming moieties. Both RosA1 and RosA2 contain six cysteine residues, with twelve and five Ser/Thr residues, respectively. Alignment of RosA peptides using MUSCLE with previously characterized two-component lantibiotics could not conclusively identify the α-peptide, but the β-component was found conserved and extended (Figure 1C and 1D). So, the putative α-peptide which displayed limited sequence similarity on the N-terminus, was designated as RosA2α, while β-peptide was designated as RosA1β. In general, α- and β-component of roseocin is extended than the ones that are previously characterized.

**Figure 1.**
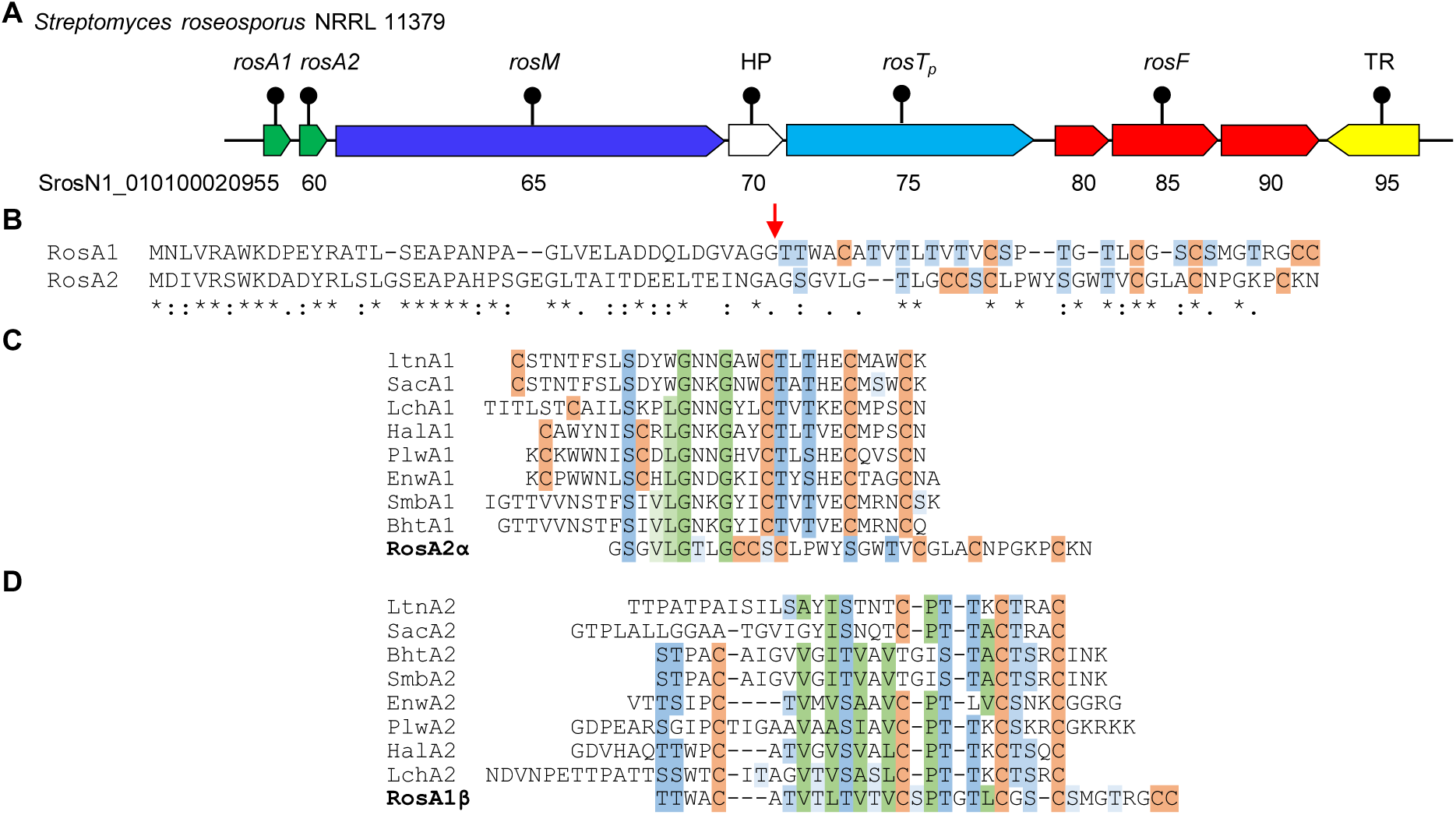
Roseocin is a homolog of two-component lantibiotics. (A) The roseocin biosynthetic gene cluster encoded in *S. roseosporus* NRRL 11379. Locus tags are mentioned below the genes. (B) Alignment of the full-length RosA1 and RosA2. The cleavage site for the C39 protease of RosT_p_ is present next to the double glycine motif (marked with an arrow). (C) & (D) Multiple sequence alignment of RosA2α (ZP_04710450.1) and RosA1β (ZP_04710449.1) core-peptides with α- and β-peptides of two-component lantibiotics lacticin 3147 (LtnA1, LtnA2); staphylococcin C55 (SacA1, SacA2); plantaricin W (PlwA1, PlwA2); BhtA (BhtA1, BhtA2); lichenicidin (LchA1, LchA2); haloduracin (HalA1, HalA2) and enterocin W (EnwA1, EnwA2). HP-hypothetical protein. See also Table S1 and S5.

### His_6_-mRosA1 and His_6_-mRosA2 are dehydrated by RosM *in vivo*

To obtain the post-translationally modified RosA peptides, two recombinant plasmids were constructed, each having lanthionine synthetase *rosM,* together with either *rosA1* (pRSFDuet-*rosA1*-*rosM*) or *rosA2* (pRSFDuet-*rosA2-rosM*). Two *E. coli* BL21(DE3) hosts, harboring each of these constructs, were induced with IPTG for expression and the peptides were found in soluble fraction. The hexahistidine tagged peptides were purified by Ni-NTA affinity chromatography followed by RP-HPLC. A yield of approximately 4 mg of HPLC purified product was obtained for each of the RosA peptides from one liter culture. As expected, the two hexahistidine tagged RosA peptides were found to be modified by RosM, as MALDI-TOF MS analysis identified a reduction of mass in multiple of 18 Da from the calculated mass of the unmodified peptides (Figure 2A and 2B). His_6_-mRosA1 (m-indicates modified) was nine-fold dehydrated at 8803 *m/z* (calc. mono. 8803 *m/z* [M-9H_2_O+Na]^+^), with the N-terminal methionine excised by the inherent activity of *E. coli* N-aminopeptidase (Lowther and Matthews, 2000). Similarly, His_6_-mRosA2 is four-fold dehydrated at 9558 *m/z* (calc. mono. 9558 *m/z* [M-4H_2_O+Na]^+^) along with the methionine excision. In both the peptides, number of dehydration were less than the total number of Ser/Thr residues present, which is not uncommon in lanthipeptide biosynthesis (Xie *et al.*, 2004), as often not all these dehydrated residues are linked into lanthionine bridges (McClerren *et al.*, 2006). An additional peak for a gluconoylated derivative was also observed for both the hexahistidine tagged RosA peptides with a mass shift of +178 Da and +258 Da. As per earlier reports, proteins that are expressed in *E. coli* with an N-terminus Gly-Ser-Ser-[His]_6_-undergo spontaneous α-N-6-phosphogluconoylation (+258 Da) and upon removal of the phosphate by the host cell phosphatase, the mass shift becomes +178 Da (Geoghegan *et al.*, 1999). However, these extra peaks for the gluconoylated product do not affect the final bioactivity, as the N-terminal region gets removed upon the *in vitro* proteolytic removal of the leader peptide. In conclusion, MALDI-TOF MS analysis confirmed the successful post-translational modification of both His_6_-mRosA1 and His_6_-mRosA2 by single RosM, heterologously. Although mass analysis of purified peptide can detect the number of dehydrations, the extent of cyclization cannot be determined as there is no change in mass upon the intra-peptide Michael addition of cysteine residues to the dehydrated residues, during lanthionine ring formation.

**Figure 2.**
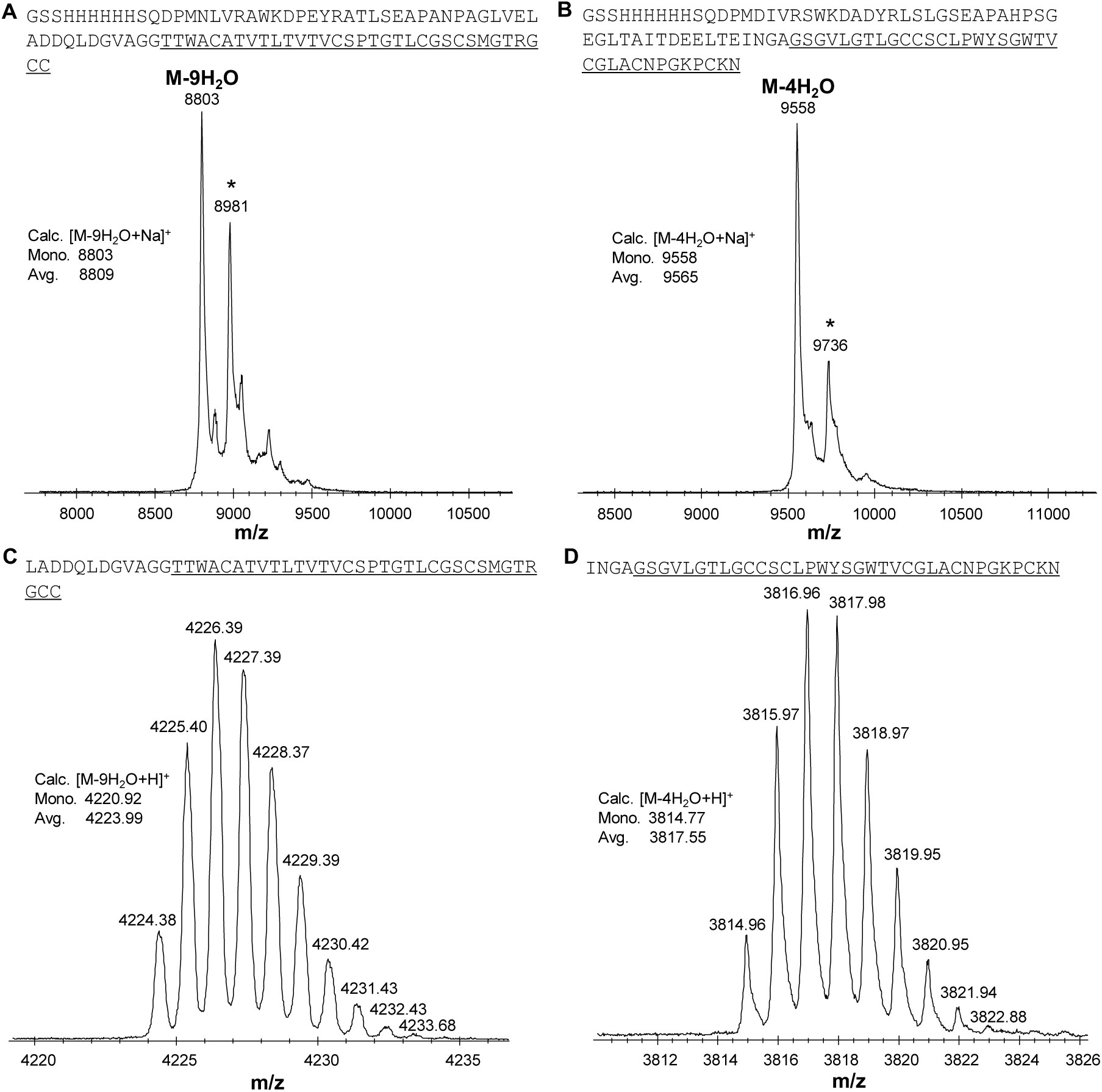
His_6_-mRosA1 and His_6_-mRosA2 are dehydrated by RosM in *E. coli.* MALDI-TOF MS spectra of purified His_6_-mRosA1 and His_6_-mRosA2. (A) His_6_-mRosA1 was nine-fold dehydrated with an observed mass of 8803 *m/z* (calc. mono. 8803 *m/z* [M-9H_2_O+Na]^+^). (B) His_6_-mRosA2 was four-fold dehydrated with an observed ma**s**s of 9558 *m/z* (calc. mono. 9558 *m/z* [M-4H_2_O+Na]^+^). An additional peak for α-N-gluconoylated product (at +178 Da) was also observed for both the heterologously produced RosA peptides (marked with an asterisk). Both His_6_-mRosA1 and His_6_-mRosA2 were treated with GluC for high-resolution mass spectrometric analysis. (C) Isotopically resolved nine-fold dehydrated RosA1β’ with 4224.38 *m/z* (calc. mono. 4220.92 *m/z* [M-9H_2_O+H]^+^) and (D) RosA2α’ with 3814.96 *m/z* (calc. mono. 3814.77 *m/z* [M-4H_2_O+H]^+^). Core peptide region is highlighted with an underline. See also Figure S1

### His_6_-mRosA1 has six, while His_6_-mRosA2 has four lanthionine rings

Cysteine residue not involved in lanthionine ring can be detected by alkylation with iodoacetamide (IAA) and observed with mass spectrometry. A mass shift of +57.05 Da is observed when a carbamidomethyl adduct with free cysteine is formed (McClerren *et al.*, 2006). So, the alkylation of modified peptides was carried out in reducing conditions followed by MALDI-TOF MS analysis. Alkylation of His_6_-mRosA1 in the presence of tris(2-carboxyethyl) phosphine (TCEP) did not lead to any change in its mass, indicating that none of the six cysteine residues are free, and are cyclized to six lanthionine rings. Alkylation of His_6_-mRosA2 in the presence of TCEP indicated an increase of mass corresponding to carbamidomethylation of two of the six cysteine residues with 9679 *m/z* (calc. avg. 9679 *m/z* [M-4H_2_O+2IAA+Na]^+^; Figure 3). In conclusion, His_6_-mRosA1 and His_6_-mRosA2 are installed with six and four lanthionine rings, respectively. The observation of two alkylations in His_6_-mRosA2, under reducing condition directed towards the possibility of a disulfide linkage, which was further investigated following the leader peptide removal.

**Figure 3.**
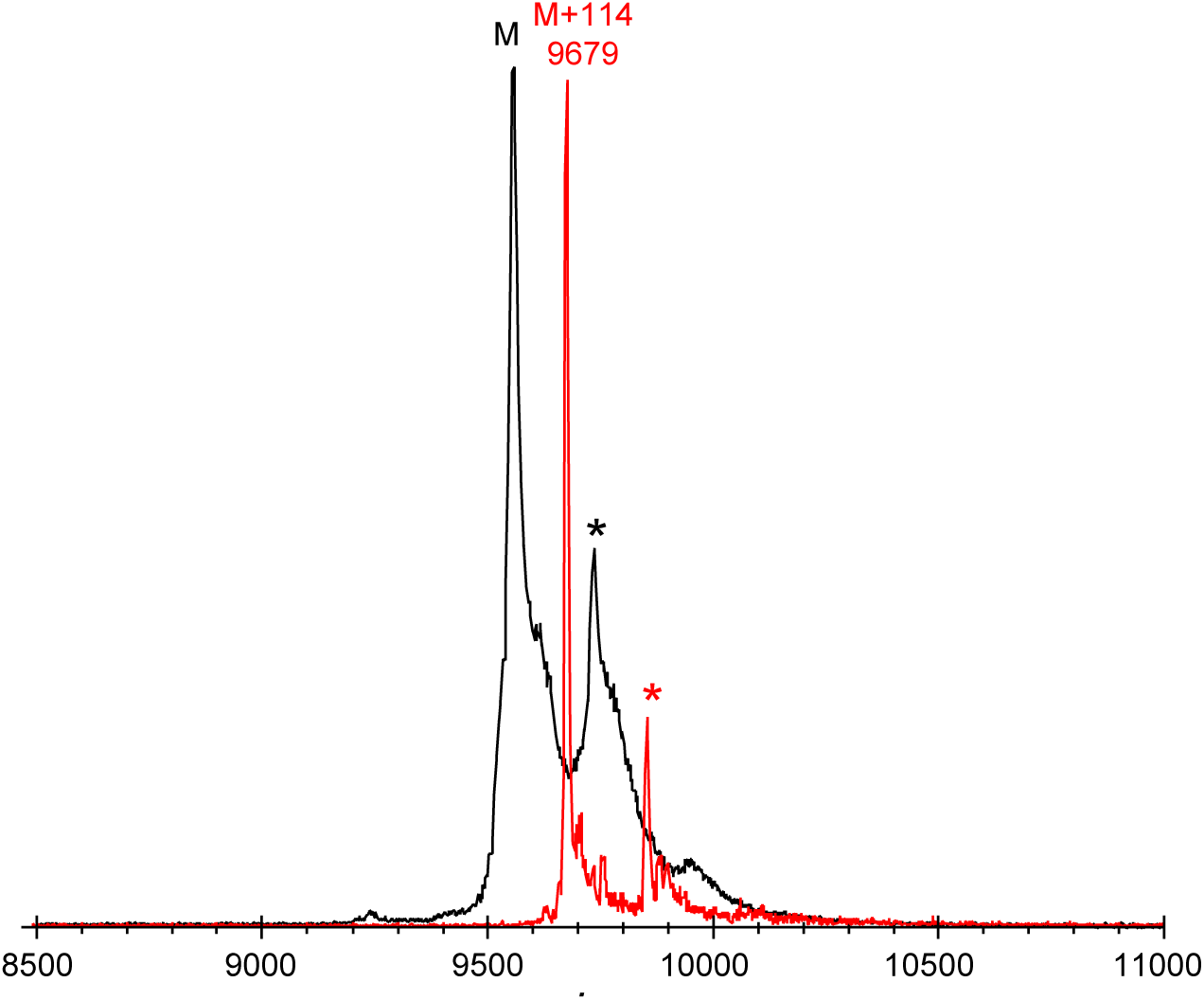
His_6_-mRosA1 has six while His_6_-mRosA2 has four lanthionine rings. Both His_6_-mRosA1 and His_6_-mRosA2 were subjected to alkylation with IAA and analyzed with MALDI TOF MS. For His_6_-mRosA2, a mass shift corresponding to two carbamidomethylation (+114 Da) was observed at 9679 *m/z* (calc. avg. 9679 *m/z* [M+Na+2IAA]^+^) indicating that two of the six cysteines residues are not cyclized. In the case of His_6_-mRosA1, no new peaks were observed upon alkylation (hence not shown) indicating that all of the six cysteine residues are cyclized. See also Figure S2

### The core-peptides RosA2α’ and RosA1β’ contain multiple overlapping thioether bridges

Multiple sequence alignment (MSA) of RosA peptides identified the conserved GA/GG motif which demarcates the boundary between the leader and the core-peptide (Figure 1B). Leader peptides ending in GA/GG motif are proteolytically removed by the N-terminal C39 cysteine protease domain of the bifunctional transporter LanT_p_ and indeed such a domain is present in RosT_p_ of the roseocin cluster (Figure 1A). For *in vitro* studies, removal of the leader peptide has been achieved using commercial proteases such as LysC, trypsin or GluC to obtain bioactive lanthipeptides (Shi *et al.*, 2011). Since, both the RosA peptides have multiple GluC cleavage sites in the leader region (at the C-terminal of glutamate residues) and none in the core-peptide, we utilized GluC for *in-vitro* removal of the leader peptide along with the hexahistidine tag. Treatment of RosA peptides with GluC removed the leader peptide, thereby reducing the size to <5kDa and thus allowed high-resolution MALDI-TOF MS, and sequence analysis with tandem MS. The GluC treated fragments were identified to be proteolytically cleaved at 3^rd^ and 5^th^ Glu residues for RosA1 and RosA2, respectively (Figure 2C and 2D). The RosA core peptides, with the four and twelve residue trace of the leader still attached were termed as RosA2α’ and RosA1β’.

In general, as reported for other lantibiotics, fragmentation was not observed in the region protected with lanthionine rings. In case of RosA2α’, for which a probable disulfide bond presence was speculated by alkylation assay, MS/MS analysis was carried out with both the reduced and non-reduced forms of peptide. An increase in mass by ∼2Da was observed upon TCEP treatment of RosA2α’, indicating an addition of two protons to the disulfide bonded Cys residues (Figure S2). Reduction of RosA2α’ also led to an enhancement in fragmentation (Figure 4A and S3) and hence identification of b and y ions with mass shifts corresponding to dehydrations of Ser/Thr residues (Table S2). MS^n^ analysis of reduced RosA2α’ identified fragment ions on either side of Cys13 and Cys33, which indicated that these regions are not protected or involved in lanthionine rings. Furthermore, it was noted that RosA2α’ underwent alkylation only upon its reduction with TCEP (Figure S2), and MS/MS analysis of the alkylated RosA2α’ also identified Cys13 and Cys33 residues to be carbamidomethylated (Figure S4). Identification of distant cysteines as disulfide partners indicated RosA2α’ to be having a conformationally constrained structure. Also, no fragment ions were observed in two long stretches indicating the presence of thioether bridges in the region (Figure 4A). For RosA1β’, MS/MS data along with the absence of alkylation, suggested cyclization of all the six cysteines into lanthionine rings (Figure 4B). Fragment ions were majorly observed for the N-terminus and a few ions in the C-terminus indicating lanthionine protection (Table S3).

**Figure 4.**
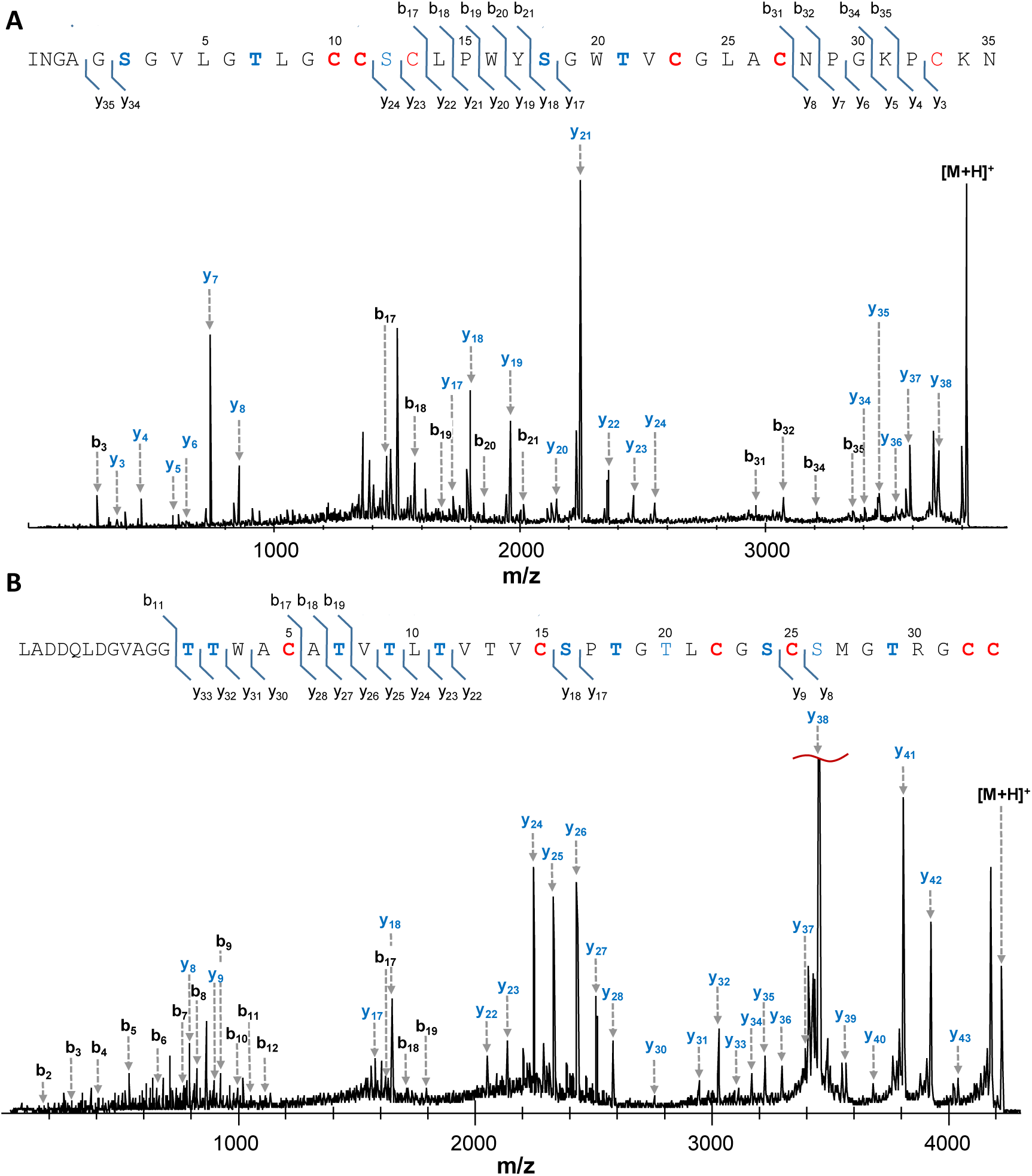
RosA2α’ and RosA1β’ core-peptides contain overlapping thioether bridges. Fragment ions were not observed in the region protected with lanthionine rings and the mass shifts corresponding to dehydrated Ser/Thr are observed. The b and y ions observed in MALDI TOF MS/MS spectrum for (A) RosA2α’ and (B) RosA1β’. Residues in bold are modified by RosM. [M+H]^+^ indicates precursor ion. See also Figure S3, and Table S2 and S3

### RosA2α’ ands RosA1β’ display synergistic antimicrobial activity

RosA peptides were tested for their antimicrobial activity against *L. monocytogenes* MTCC 839, *Bacillus subtilis* MTCC 121, *Staphylococcus aureus* MTCC 1430. *Pseudomonas aeruginosa* MTCC 1934, *Escherichia coli* MTCC 1610 and *Vibrio cholerae* MTCC 3904. RosA2α’ and RosA1β’ did not display any antimicrobial activity when tested individually, but a zone of inhibition was observed against Gram-positive bacteria when both the peptides were spotted together (Table 1). Uncleaved His_6_-mRosA1 and His_6_-mRosA2 peptides alone and in combination did not display any antimicrobial activity, as was expected from a leader peptide attached lantibiotic (Figure 5). Such a synergistic antimicrobial activity of two separate post-translationally modified peptides, which display little to no activity alone is a characteristic of two-component lantibiotics (Navaratna et al., 1998). For roseocin, antimicrobial activity was observed against tested Gram-positive bacteria except for *S. aureus* MTCC 1430 and weak to no activity was observed against Gram-negative bacteria. Another exception to this was the observation of a zone of inhibition with RosA2α’ alone, against *B. subtilis* MTCC 121, which enhanced to a distinct zone with the addition of RosA1β’ (not shown).

**Table 1.** Bioactivity analysis of roseocin against Gram-positive and Gram-negative bacteria. Roseocin displayed synergistic antimicrobial activity against Gram-positive bacteria, and weak to no activity against Gram-negative bacteria.

| Indicator strain | RosA2α | RosA1α | Combined |
| --- | --- | --- | --- |
| **Gram-positive bacteria** |  |  |  |
| *Listeria monocytogenes* MTCC 839 | - | - | +++ |
| *Bacillus subtilis* MTCC 121 | ++ | - | +++ |
| *Staphylococcus aureus* MTCC 1430 | - | - | - |
| **Gram-negative bacteria** |  |  |  |
| *Pseudomonas aeruginosa* MTCC 1934 | - | - | - |
| *Escherichia coli* MTCC 1610 | - | - | + |
| *Vibrio cholerae* MTCC 3904 | - | - | - |

**Figure 5.**
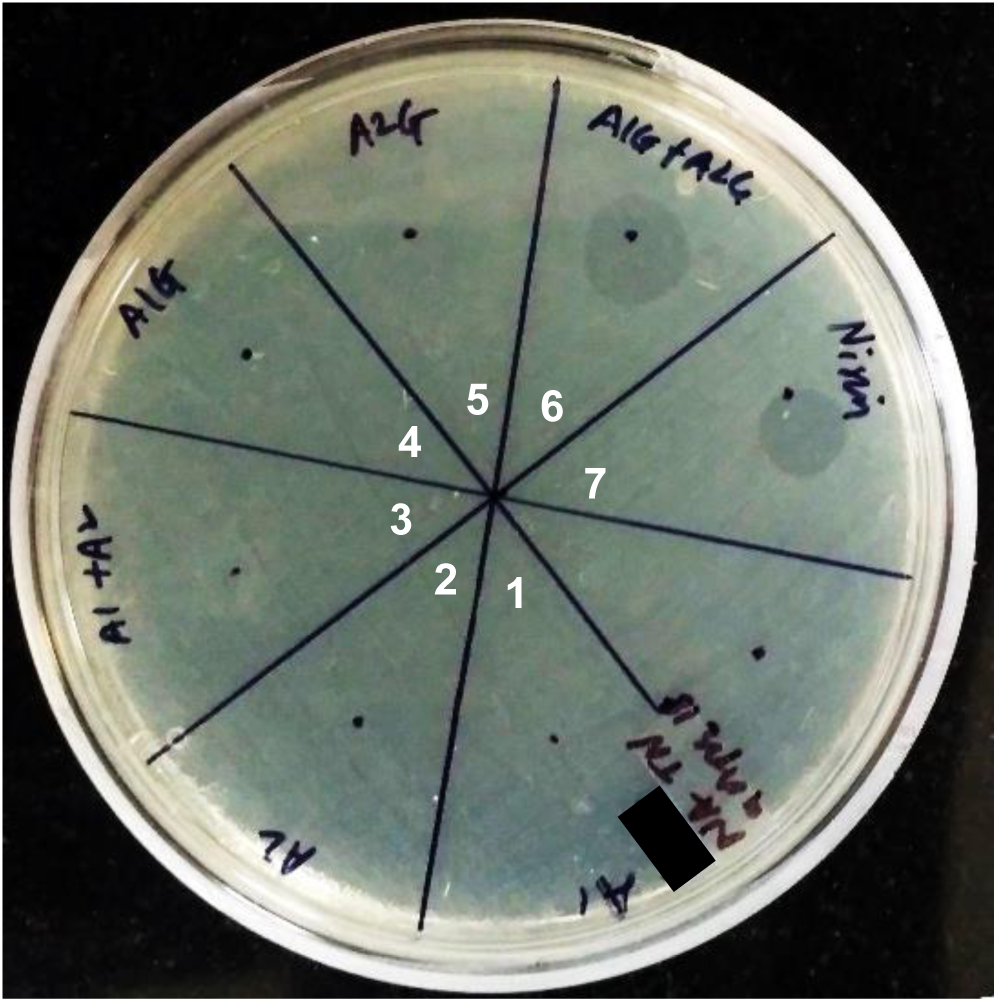
RosA2α’ and RosA1β’ displayed synergistic antimicrobial activity against *L. monocytogenes* 839. A zone of inhibition was observed only when both the leader removed core peptides were spotted together. RosA peptides were tested at 30 µM concentration. Sector-1, 40 µL His_6_-mRosA1; Sector-2, 40 µL His_6_-mRosA2; Sector-3, combined His_6_-mRosA1 (20 µL) and His_6_-mRosA2 (20 µL); Sector-4, 40 µL GluC digest of His_6_-mRosA2 (to generate RosA2α’); Sector-5, 40 µL GluC digest of His_6_-RosA1 (to generate RosA1β’); Sector-5, combined digests (20 µL RosA2α’ and 20 µL RosA1β’); Sector-7, 3 µL of 300 µM nisin.

Extent of antimicrobial activity indicated by: (-) no detectable activity; (+) weak; (++) moderate; (+++) high activity.

## DISCUSSION

Lantibiotics are believed to be the solution for the current problem of antimicrobial resistance and hence researchers are discovering novel lantibiotics through various approaches. The earlier approach was dependent upon culture-based activity screening with the limitation of re-discovery of known compounds, being time-consuming and is also subject to the selective conditions in which the culture can be induced for the secondary metabolite production. However, the currently followed *in silico* approach involves the identification of potential producers which can then be confidently taken up for wet lab experiments. In our initial attempt to identify class II lantibiotic biosynthetic gene clusters encoded in the sequenced bacteria, we reported several novel putative lantibiotic clusters. Two of these clusters reported in our study have already been characterized to encode two-component lantibiotics, bicereucin and flavecins (Huo and van der Donk, 2016; Zhao and van der Donk, 2016). The lantibiotic cluster characterized in the current study attracted our attention for its special attributes like a single RosM for processing of two RosA precursor peptides which are homologs of two-LanM processed two-component lantibiotics (Figure 1). These peptides have altogether different core region, extended C-terminus and excessive Ser/Thr and Cys residues. The putative two-component lantibiotic from this *Actinomycete* was expected to be having more lanthionine rings than already reported lantibiotics, isolated from *Firmicutes* and hence would probably be more efficacious.

In a previous study, to interrogate the biosynthetic capacity of the potential producers, Kersten *et al*. (2011) executed a MS-guided genome mining approach for natural product peptidogenomics in eight different *Streptomyces* strains, which are the known producers of various antimicrobial natural products (Kersten *et al.*, 2011). The enormous MS data of *Streptomyces roseosporus* NRRL 15998 along with the *in silico* analysis of the genome provided by them mention the presence of a similar cluster with two precursor peptide (73 aa and 80 aa, respectively), without the identification of immunity genes. Moreover, this previous study didn’t focus on the bioactivity analysis of the lanthipeptides. Also, the product of only one of the two LanA genes could be detected by them from the n-butanolic extract of *S. roseosporus,* and in the absence of the second gene product, it could not be studied either. However, our successful heterologous production of the two modified RosA peptides by *in vivo* promiscuous activity of RosM, followed by *in-vitro* leader peptide removal for generation of bioactive lanthipeptides led to the structure prediction and bioactivity evaluation of the encoded two-component lantibiotic roseocin.

The structure of RosA2α, as suggested by sequence homology and tandem MS analysis, was unlike any other previously known α-component. It was initially speculated that with six cysteine residues, the RosA2α’ would be installed with only five lanthionine rings due to the availability of only five Ser/Thr residues, leaving one of the six cysteine residues uncylized (Figure 1C). Unexpectedly, MALDI TOF MS analysis of the RosA2α’ confirmed only four dehydrations (Figure 2B and 2D), further limiting the number of expected lanthionine rings from five to only four, leaving two of the six cysteine residues uncyclized and one of the Ser/Thr undehydrated. These unmodified residues were identified by sequence analysis with tandem MS, both with, and without the iodoacetamide treatment of TCEP reduced RosA2α’. The residues Cys13, Cys33 and Ser12 were found to be unmodified by RosM (Figure 4A and S4). Moreover, alkylation was observed only in the presence of a reducing agent which indicated that these residues are involved in a disulfide bridge and hence are in the oxidized state naturally (Figure S2). While other two-component lantibiotics like Halα, Plwα, and Enwα have a disulfide bond between the two nearby cysteine residues, in RosA2α’ these cysteine residues are distant and are found to be flanking two thioether bridges (Figure 6). The presence of this disulfide bond in RosA2α’ contributed to a globular structure and prevented fragmentation in MS/MS for sequence analysis, which was made possible by reduction of this disulfide bond (Figure S3). Interestingly, the conserved lipid II binding CTxTxEC motif found in the α-component of all the haloduracin related two-component lantibiotics, is absent in RosA2α’ peptide. It is perplexing in light of its importance for interaction with lipid II and its conservation in all the α-peptides characterized till date.

**Figure 6.**
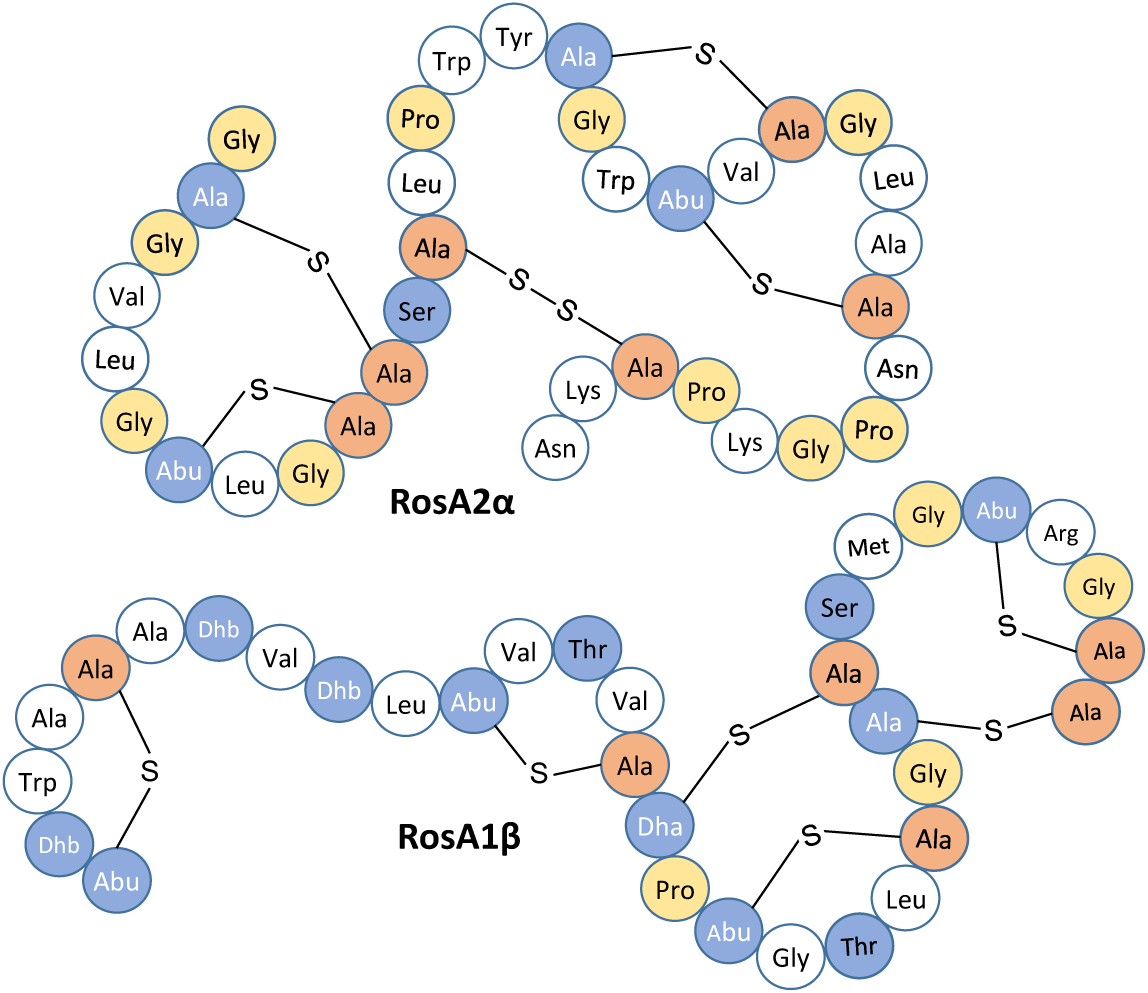
Structure of RosA2α and RosA1β. Predicted structures of RosA2α and RosA1β as per the fragmentation and mass shifts obtained in the tandem MS analysis. RosA2α contains a disulfide bond between Cys13 and Cys33 that flank the two of the four lanthionine rings. RosA1β is installed with six lanthionine rings and its structure was predicted using Halβ as the template, tandem MS analysis, and alkylation assay.

RosA1β’ with twelve Ser/Thr was speculated to undergo sufficient dehydration to form lanthionine rings with six cysteines. The mass analysis confirmed nine dehydrations of the total twelve residues (Figure 2A and 2C), which were still sufficient for six cysteines to undergo Michael addition to these residues for lanthionine formation. Indeed, the presence of six lanthionine rings was confirmed by the absence of alkylation upon iodoacetamide treatment. The structure of RosA1β’, as suggested by sequence homology and tandem MS data, is consistent with those of the other β-peptides for at least first three rings on the N-terminus (Figure 1D and 4B). Besides being an elongated peptide, RosA1β’ contains two additional lanthionine rings than the usual four in β-peptides of the other two-component lantibiotics. The presence of extra lanthionine rings could endow the peptide with a more constrained and stable structure (Figure 6), and the length extension might allow better spanning of the bacterial membrane. As with the presence of multiple sites for Michael addition (nine dehydrated residues for six cysteines), RosA1β’ could undertake myriad of configurations. Lack of fragmentation from the middle of the peptide till the C-terminus, probably due to thioether bridge overlaps made the assignment of the ring structure in this region difficult (Figure 4B). Additionally, as the C-terminal is extended and does not show homology with known lantibiotics, a conclusive rings pattern could only be predicted. The methyllanthionine rings encoded by the Dhx-Dhx-Xxx-Xxx-Cys motif at the N-terminus of RosA1β, possibly has LL stereochemistry, instead of the usual DL stereochemistry. This motif has previously been associated with the formation of LL stereochemistry in flavecins (Zhao and van der Donk, 2016) and carnolysin (Lohans *et al.*, 2014).

As with the other two-component lantibiotics, roseocin displayed synergistic antimicrobial activity primarily against the Gram-positive bacteria (Table 1) and only weak activity if any against the Gram-negative bacteria. The bioactivity was obtained even with the four- and twelve-residue long traces of the leader peptides still attached to RosA2α’ and RosA1β’, respectively (Figure 5). The extent of proteolytic processing of lanthipeptides can be a determining factor for the bioactivity and hence, it is speculated that an enhancement would be observed upon removal of the remaining trace residues. In case of the class I lantibiotic gallidermin, a leader trace of similar length (12 aa) prevented any antimicrobial activity and removal of this trace was necessary for it to be antimicrobial (Valsesia *et al.*, 2007). The type II, two-component lantibiotic lichenicidin is first processed by a bifunctional LicT_p_ for removal of the leader peptide, which is followed by the removal of a small oligopeptide (6 aa) by subtilisin family protease LicP in order to be bioactive (Caetano *et al.*, 2011). Previous studies, where a commercial protease was utilized for leader peptide cleavage, relied on N-aminopeptidase for removal of the trace left behind by GluC treatment (Garg *et al.*, 2012; Zhao and van der Donk, 2016). This strategy was hindered in case of RosA peptides, as N-aminopeptidase activity is obstructed by the aspartate residues present in the leader trace of RosA1β’. As the trace allowed bioactivity and structural analysis, it was not deemed necessary to remove these residues in a second step. In future, we aim to co-express RosT_p_ in a compatible plasmid for *in vivo* cleavage of the leader peptide, thus leading to the extracellular transport of the fully processed native roseocin peptides - RosA2α, and RosA1β (Wang *et al.*, 2016), to ascertain any alteration or enhancement in bioactivity and for further mechanistic insights. However, the purification strategy will have to be worked-out accordingly because of the absence of the hexahistidine tag. The spectrum of Gram-positive bacteria inhibited by roseocin is required to be screened further, and the weak bioactivity against Gram-negative bacteria is ought to be enhanced by bioengineering or using hurdle technology as observed for the lantibiotic nisin (Field *et al.*, 2012).

A recent study described large-scale analysis of lanthipeptide biosynthetic gene clusters from *Actinobacteria* and grouped these clusters into gene cluster families (GCF) (Zhang *et al.*, 2015). Putative two-component lantibiotics with twelve members were grouped in Lant_GCF.30 (encoding two LanM and two LanA substrates), none of which have been characterized experimentally. Additionally, the currently known few lantibiotics reported from *Actinobacteria* are all single peptide lantibiotics. Here, we characterized a Lant_GCF.73 group member as a two-component lantibiotic from *S. roseosporus* NRRL 11379, which comprises of a single RosM for the synthesis of RosA2α and RosA1β. The alpha peptide is installed with four lanthionines, and a disulfide bridge, while the beta-component comprised of six thioether bridges. Looking for the homologous clusters, we identified a biosynthetic gene cluster encoding a natural variant of roseocin in *Glycomyces harbinensis* (Labeda *et al.*, 1985). A study on such variants is important to understand the role of modifications that affect bioactivity, for designing semisynthetic derivatives with improved characteristics (Gomes *et al.*, 2017), which we plan to carry out in future. Overall, roseocin is the first example of a two-component lantibiotic from a non-*Firmicute*.

## METHODS

### CONTACT FOR REAGENT AND RESOURCE SHARING

Further information and requests for resources and reagents should be directed to and will be fulfilled by the Lead Contact, Dipti Sareen.

### EXPERIMENTAL MODEL AND SUBJECT DETAILS

*Streptomyces roseosporus* NRRL 11379 was grown in ISP2 medium (1 liter media with 4 g yeast extract, 10 g malt extract and 4 g dextrose at pH 7) at 200 rpm, 28°C for genomic DNA isolation. All the *E. coli* strains were grown in LB at 37°C, 200 rpm with or without kanamycin, as required. Indicator strains were cultured in nutrient broth at 37°C, 200 rpm, except for *B. subtilis* which was cultured at 30°C. Ready-to-use and individual media components were purchased from Himedia.

## METHOD DETAILS

### Cloning of *rosA1* and *rosA2* with *rosM* in pRSFDuet1 vector

Genomic DNA was isolated using a previously described method (Kumar *et al.*, 2011) from *S. roseosporus* NRRL 11379. The three genes were PCR amplified using respective primers with restriction sites, on a *BIO-RAD* MyCycler™ Thermal Cycler using Q5 DNA polymerase. Gene for RosA1 and RosA2 was inserted in MCS-1 of pRSFDuet-1 using BamHI and HindIII sites, and RosM in MCS-2 using NdeI and XhoI restriction sites. Chemically competent *E. coli* DH10B was prepared, transformed by heat shock and selected on a kanamycin plate. Colonies carrying recombinant plasmid (pRSFDuet-*rosA1-rosM* and pRSFDuet-*rosA2-rosM*) were screened by colony PCR, which was followed by gene sequencing of the complete ORF from SciGenome labs using appropriate vector-specific primers (sequencing primers in **Table S4**). The 3315 bp long *rosM* was additionally sequenced using gene-specific primers (M1, M2, and M3). Sequencing results were analyzed on FinchTV and gene sequence was found 100% identical to the sequence on NCBI. These constructs were then transformed into *E. coli* BL21(DE3) for production of modified RosA peptides.

### Production and purification of *in vivo* modified RosA peptides

A single colony was inoculated in 10 mL LB for overnight. The culture was used to inoculate 2-L of LB added with 50 μg/mL of kanamycin. The culture was incubated at 37°C, 200 rpm until the A_600_ reached 0.6-0.8, and the temperature was lowered at this point to 18°C and induced with 0.1 mM IPTG for an additional 24 hrs. The cells were harvested by centrifugation at 5000 g for 15 min. The cell pellet was resuspended in 50 mL start buffer (50 mM Tris-HCl, pH 8.0, 500 mM NaCl, 1 mM Imidazole, 1 mM PMSF), and cell lysis was carried out using Sonics VC 505 sonicator. The lysate was cleared by centrifugation at 25,000 g for 30 min at 4°C (on *SIGMA 3K30* using 12158 rotor). The cleared lysate was then loaded on a manually packed 2 mL Ni Sepharose™ High Performance (GE Healthcare) metal affinity resin. The resin was washed with wash buffer (start buffer with 30 mM imidazole), and the peptide was eluted in elution buffer (wash buffer containing 500 mM imidazole). Eluted fractions were analyzed on 16%/6M urea tricine SDS-PAGE (Schägger H, 2006). Further purification and desalting of the eluted fractions were done with an Agilent 300 SB-C18 semi-preparative column on an Agilent 1260 infinity series HPLC system. Sample loading and desalting was done with mobile phase A (5:95 ACN:H_2_O, with 0.1% TFA) and gradient of 0-60% of solvent B (95:5 ACN:H_2_O, with 0.085% TFA) at 4 mL/min was used to elute the peptides. Elution was monitored at 220 and 280 nm. Modified His_6_-mRosA1and His_6_-mRosA2 started eluting out at 43% and 40% of mobile phase B, respectively. Collected fractions were lyophilized and re-dissolved in MilliQ water for further analysis.

### Molecular weight and sequence analysis of RosA peptides

Matrix Assisted Laser Desorption/Ionization time-of-flight mass spectrometry (MALDI-TOF MS) was carried for determining accurate masses and post-translational modifications in the peptides, on a Bruker ultrafleXtreme™ MALDI-TOF/TOF system maintained at CSIC, PGIMER, Chandigarh. Intact peptides with mass of >5 kDa were analyzed in linear mode and proteolytic digests (peptides of <5 kDa) were analyzed in the reflectron mode (high resolution mode which leads to isotopic separation). HPLC purified peptides (in 50:50 ACN:H_2_O 0.1% TFA) were mixed with sinapinic acid and alpha-Cyano-4-hydroxycinnamic acid (5 mg/mL in 50:50 ACN:H_2_O, 0.1% TFA) for analysis in linear and reflectron mode, respectively. Mass spectra were recorded in positive ion mode. The Bruker flexControl was used for data acquisition, and SeeMS and mMass program were used for data analysis.

### Reduction and IAA modification of the RosA peptides

The purified peptides in 50 mM tris HCl pH 8.0 were (1) incubated with 1 mM TCEP at 37°C for 30 min for reduction of the disulphide bond, (2) followed by addition of 10 mM IAA for 60 min (kept in dark). The sample was processed to remove unbound IAA and/or TCEP with Pierce™ C18 spin column before analysis with MALDI-TOF MS.

### Preparation of bioactive roseocin and growth inhibition assay

Purified peptides were dissolved in sterile MilliQ grade water and treated with 0.1 μg/μl endoproteinase GluC (P8100, NEB) in 20:1 ratio at a final concentration of 30 µM of each peptides for 16 h at 37°C. The His_6_-mRosA proteolytic digests were directly used for antimicrobial activity analysis. The His_6_-mRosA1 and His_6_-mRosA2 peptides were assayed as negative controls at a final concentration of 30 uM. A 40 µl volume was used for the spot of a single peptide, while 20 µl of each was used for spots containing both the peptides; nisin was assayed using 3 µl of a 300 µM solution. Indicator strains were grown to 0.1 OD_600_ before spreading on agar plates using sterile cotton swab which was followed by spotting of the peptide mixture in 10 µL aliquots and allowed to dry before re-spotting again till the required volume is spotted. The plates were incubated for overnight at required temperature. A 300 µM stock of nisin was prepared using 2.5% nisin powder (Sigma - N5764) by dissolving 40 mg/mL in 0.05% acetic acid. It was allowed to dissolve for 10 min and then spun to remove any insoluble material.

## QUANTIFICATION AND STATISTICAL ANALYSIS

His_6_-mRosA peptides (∼10 kDa) were quantitated by densitometry using lysozyme (14.4 kDa) as a standard with ImageJ 1.48v software.

## KEY RESOURCES

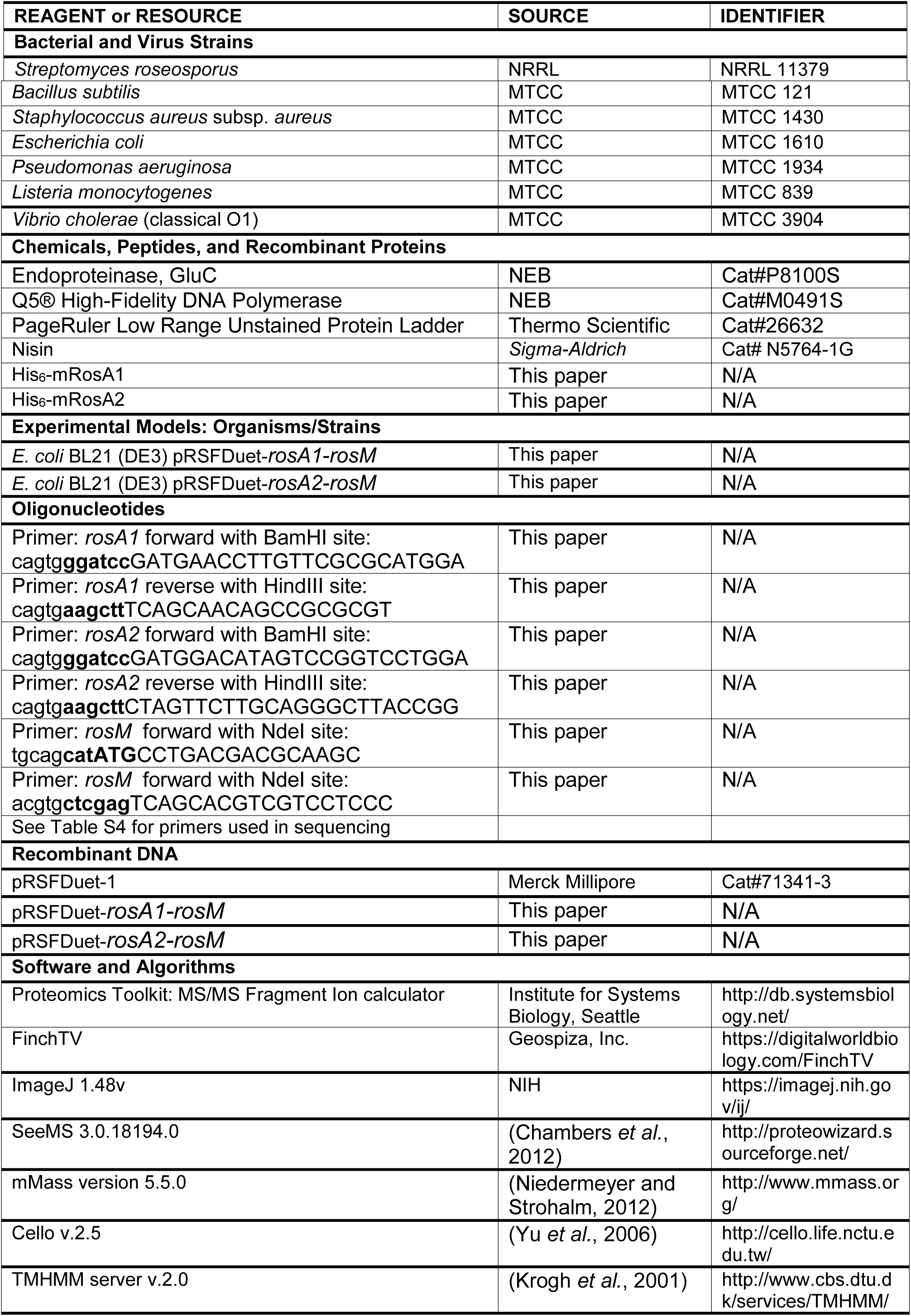

## Supporting information

## SUPPLEMENTAL INFORMATION TITLES AND LEGENDS

Supplemental information includes 4 figures, and 5 tables.

## AUTHOR CONTRIBUTIONS

Supervision D.S.; Resources D.S. Conceptualization D.S. and M.S.; Methodology D.S. and M.S.; Investigation M.S.; Formal analysis M.S.; Writing-original draft, M.S.; Writing-Review & Editing, M.S and D.S. Funding acquisition M.S. and D.S.

## ACKNOWLEDGMENTS

MS acknowledges the independent Senior Research Fellowship (SRF) No. 3/1/3/JRF-2011/HRD-99(11005), awarded by the Indian Council of Medical Research, Government of India, New Delhi. The financial assistance received to the lab from DST-PURSE and UGC-SAP grant, Government of India, New Delhi, is also acknowledged. MALDI-TOF MS facility maintained at CSIC, PGIMER, Chandigarh is acknowledged.

