## Supplementary material for "Semi-*in vitro* Reconstitution of Roseocin, a Two-Component Lantibiotic from an *Actinomycete*"

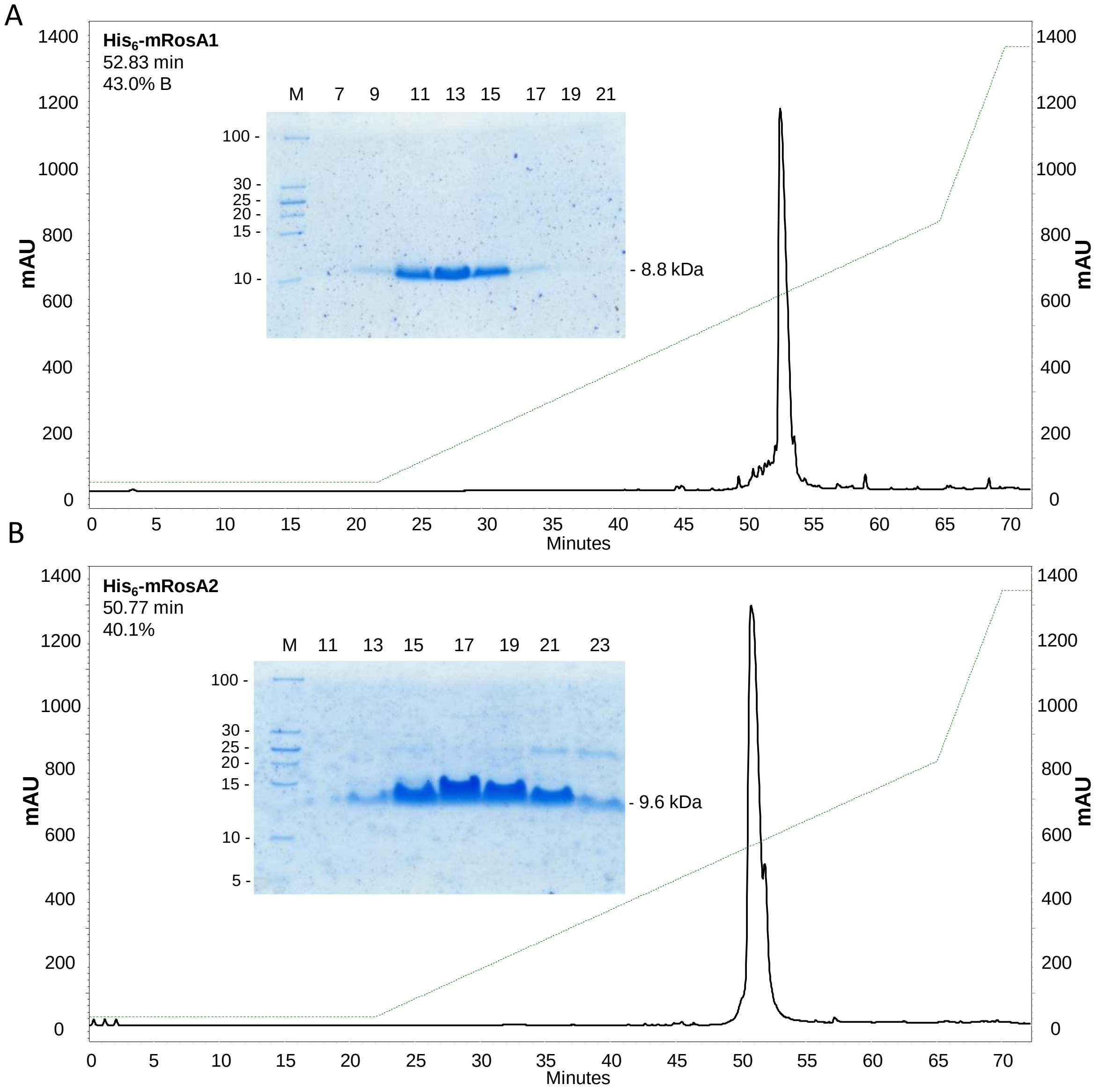


**Figure S1. Related to Figure 2. Purification of hexahistidine tagged RosA peptides with RP-HPLC.** RP-HPLC chromatogram for purification of (A) $His_6$-mRosA1 and (B) $His_6$-mRosA2. Collected fractions were lyophilized and reconstituted in MilliQ, followed by analysis on tricine SDS-PAGE (inset). Lane 1 in both the gels is low MW protein ladder and rest of the lanes are collected fractions. The spectra was noted at 280 nm.

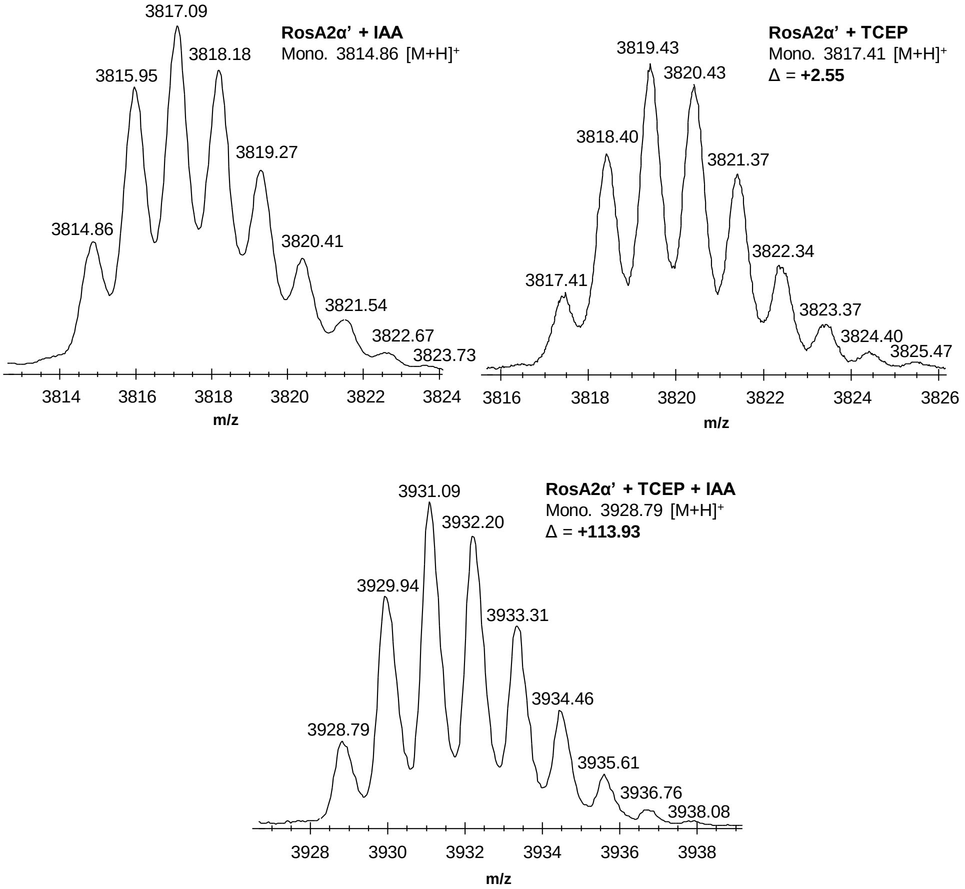


**Figure S2. Related to Figure 3. Alkylation assay of RosA2α' and analysis with high-resolution MALDI TOF MS.** $His_6$-mRosA2 was subjected to GluC digestion and given treatment with IAA only, TCEP only, and TCEP followed by IAA. (A) RosA2α' treated with IAA alone did not lead to any change in mass. (B) RosA2α' when treated with TCEP led to an increase of mass by ~2 Da. (C) Two carbamidomethylations (~ +114 Da) were observed upon reduction with TCEP followed by IAA treatment.

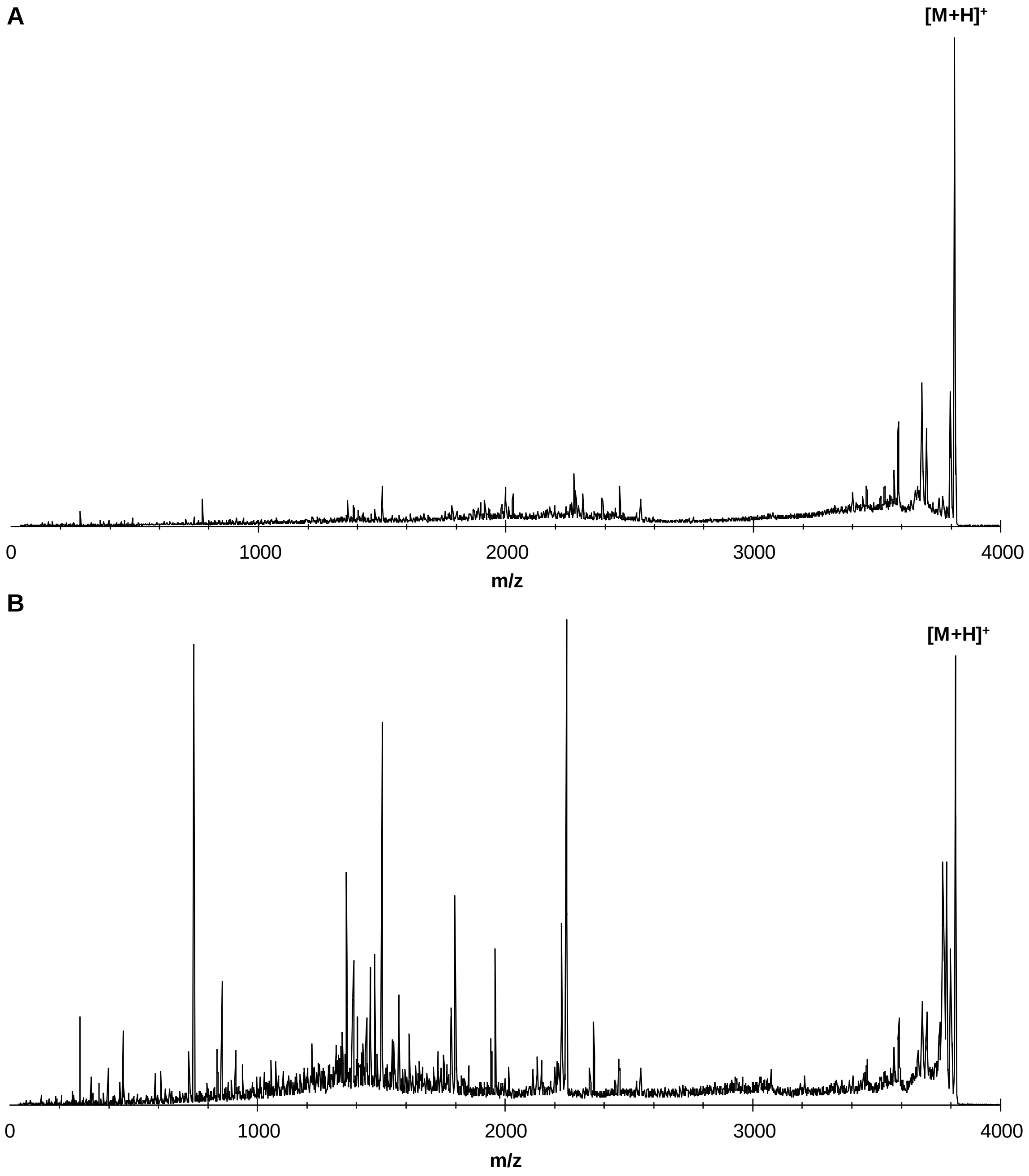


**Figure S3. Related to Figure 4. Tandem MS analysis of RosA2α' with and without TCEP treatment.** (A) Presence of disulfide bond prevented fragmentation. (B) Reduction with 1 mM TCEP led to the enhancement in fragmentation.

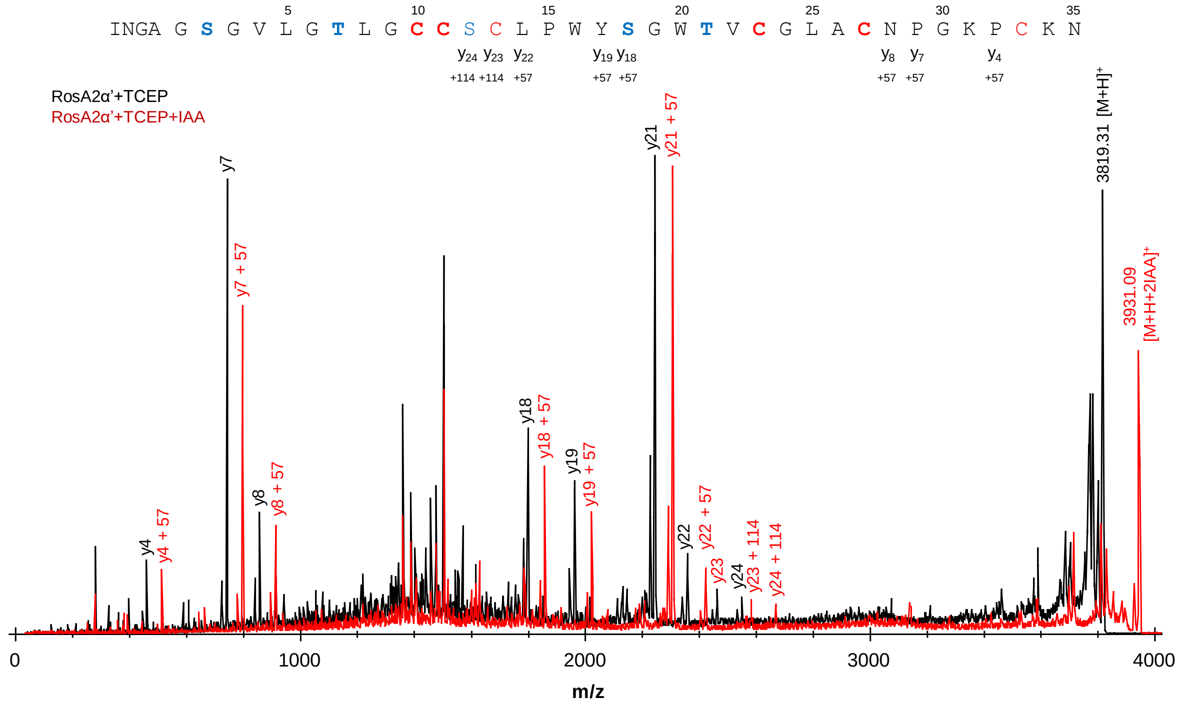


**Figure S4. Related to Figure 3. Tandem MS analysis of alkylated and non-alkylated RosA2α'.** Selective y ions are labelled for indicating the position of alkylation. Fragment ion $y_4$ to $y_{22}$ indicate one carbamidomethylation at Cys33 and $y_{23}$ indicate another at Cys13. Residues in bold are modified by RosM.

**Table S1. Bioinformatic analysis of roseocin biosynthetic gene cluster**

| **Locus tag/Accession** | **Cello v. 2.5** | **TMHMM Server 2.0** | **CDD analysis** | **Predicted function** |
|---|---|---|---|---|
| SrosN1_010100020955/ ZP_04710449.1 | Extracellular | 0 | TIGR03897: Type 2 lantibiotic, mersacidin/lichenicidin family | *rosA1* |
| SrosN1_010100020960/ ZP_04710450.1 | Extracellular | 0 | TIGR03897: Type 2 lantibiotic, mersacidin/lichenicidin family | *rosA2* |
| SrosN1_010100020965/ ZP_04710451.1 | Cytoplasmic | 0 | TIGR03897: type 2 lantibiotic biosynthesis protein LanM | *rosM* |
| SrosN1_010100020970/ ZP_04710452.1 | Cytoplasmic | 1 | No putative conserved domains | Hypothetical protein |
| SrosN1_010100020975/ ZP_04710452.1 | Membrane | 6 | COG2274: ABC-type bacteriocin/lantibiotic exporters, contain an N-terminal double-glycine peptidase domain | *rosT* |
| SrosN1_010100020980/ ZP_04710454.1 | Membrane | 4 | No conserved domains, 94% identity with Multidrug transporter (*Streptomyces sp.* Ncost-T6T-1) | Immunity* |
| SrosN1_010100020985/ ZP_04710455.1 | Cytoplasmic | 0 | cd03268: ATP-binding cassette domain of the bacitracin-resistance transporter | *rosF* |
| SrosN1_010100020990/ ZP_04710456.1 | Membrane | 6 | No conserved domains, | Immunity* |
| SrosN1_010100020995/ ZP_04710457.1 | Cytoplasmic | 0 | smart00862: Transcriptional regulatory protein - C terminal, and pfam03704: Bacterial transcriptional activator domain with TPR repeats. | Transcriptional regulator |

*The immunity genes are often found encoded in vicinity and the encoded proteins were predicted as transmembrane proteins by Cello and TMHMM servers, with multiple transmembrane helices.

**Table S2. b and y ions observed for RosA2α' in MALDI TOF MS/MS analysis (related to figure 4A)**

| Species | Calc. mass [Da] | Obs. mass [Da] | Difference [Da] | Error |
|---|---|---|---|---|
| $b_3$ | 285.156 | 285.830 | 0.674 | 0.002364 |
| $y_3$ | 364.165 | 364.210 | 0.045 | 0.000124 |
| $y_4$ | 461.218 | 462.231 | 1.013 | 0.002196 |
| $y_5$ | 589.313 | 590.326 | 1.013 | 0.001719 |
| $y_6$ | 646.334 | 647.359 | 1.025 | 0.001586 |
| $y_7$ | 743.387 | 744.412 | 1.025 | 0.001379 |
| $y_8$ | 857.423 | 858.458 | 1.028 | 0.001199 |
| $b_{17}$-$2H_2O$ | 1457.654 | 1459.668 | 2.013 | 0.001381 |
| $b_{18}$-$2H_2O$ | 1570.739 | 1572.752 | 2.013 | 0.001282 |
| $b_{19}$-$2H_2O$ | 1667.791 | 1669.805 | 2.014 | 0.001208 |
| $y_{17}$-$1H_2O$ | 1729.808 | 1731.058 | 1.250 | 0.000723 |
| $y_{18}$-$2H_2O$ | 1798.84 | 1799.842 | 1.002 | 0.000557 |
| $b_{20}$-$2H_2O$ | 1853.871 | 1855.884 | 2.013 | 0.001086 |
| $y_{19}$-$2H_2O$ | 1961.903 | 1962.905 | 1.002 | 0.000511 |
| $b_{21}$-$2H_2O$ | 2016.934 | 2018.947 | 2.013 | 0.000998 |
| $y_{20}$-$2H_2O$ | 2147.982 | 2148.985 | 1.003 | 0.000467 |
| $y_{21}$-$2H_2O$ | 2245.035 | 2246.037 | 1.002 | 0.000446 |
| $y_{22}$-$2H_2O$ | 2358.119 | 2359.121 | 1.002 | 0.000425 |
| $y_{23}$-$2H_2O$ | 2461.128 | 2463.267 | 2.137 | 0.000875 |
| $y_{24}$-$2H_2O$ | 2548.160 | 2550.390 | 2.230 | 0.000875 |
| $b_{31}$-$4H_2O$ | 2958.344 | 2960.489 | 2.145 | 0.000725 |
| $b_{32}$-$4H_2O$ | 3072.387 | 3074.592 | 2.205 | 0.000718 |
| $b_{34}$-$4H_2O$ | 3226.461 | 3228.463 | 2.002 | 0.000620 |
| $b_{35}$-$4H_2O$ | 3354.556 | 3356.558 | 2.002 | 0.000597 |
| $y_{34}$-$4H_2O$ | 3402.559 | 3405.040 | 2.481 | 0.000729 |
| $y_{35}$-$4H_2O$ | 3459.581 | 3462.091 | 2.510 | 0.000726 |
| $y_{36}$-$4H_2O$ | 3530.618 | 3533.169 | 2.551 | 0.000723 |
| $y_{37}$-$4H_2O$ | 3587.640 | 3590.221 | 2.582 | 0.000720 |
| $y_{38}$-$4H_2O$ | 3701.682 | 3704.323 | 2.641 | 0.000713 |

**Table S3. b and y ions observed for RosA1β' in MALDI TOF MS/MS analysis (related to figure 4B)**

| Species | Calc. [Da] | Obs. [Da] | Difference [Da] | Error |
|---|---|---|---|---|
| $b_2$ | **185.128** | **185.212** | **0.083** | **0.000448** |
| $b_3$ | 300.155 | 301.130 | 0.974 | 0.003245 |
| $b_4$ | 415.182 | 416.156 | 0.974 | 0.002346 |
| $b_5$ | 543.241 | 544.215 | 0.974 | 0.001793 |
| $b_6$ | 656.325 | 657.299 | 0.974 | 0.001484 |
| $b_7$ | 771.352 | 771.794 | 0.442 | 0.000573 |
| $y_8$ | 796.301 | 796.476 | 0.175 | 0.000220 |
| $b_8$ | 828.373 | 828.845 | 0.472 | 0.000569 |
| $y_9$-1$H_2O$ | 899.310 | 900.238 | 0.928 | 0.001031 |
| $b_9$ | 927.442 | 928.416 | 0.974 | 0.001050 |
| $b_{10}$ | 998.479 | 999.453 | 0.974 | 0.000975 |
| $b_{11}$ | 1055.500 | 1056.474 | 0.973 | 0.000922 |
| $b_{12}$ | 1112.522 | 1113.157 | 0.635 | 0.000571 |
| $y_{17}$-3$H_2O$ | 1579.626 | 1580.883 | 1.257 | 0.000796 |
| $b_{17}$-2$H_2O$ | 1638.743 | 1639.768 | 1.025 | 0.000612 |
| $y_{18}$-4$H_2O$ | 1648.658 | 1649.865 | 1.207 | 0.000732 |
| $b_{18}$-2$H_2O$ | 1709.780 | 1710.846 | 1.066 | 0.000610 |
| $b_{19}$-3$H_2O$ | 1792.828 | 1793.934 | 1.107 | 0.000599 |
| $y_{22}$-4$H_2O$ | 2050.852 | 2052.456 | 1.604 | 0.000782 |
| $y_{23}$-5$H_2O$ | 2133.899 | 2136.106 | 2.207 | 0.001043 |
| $y_{24}$-5$H_2O$ | 2246.983 | 2249.190 | 2.207 | 0.000990 |
| $y_{25}$-6$H_2O$ | 2330.031 | 2332.238 | 2.207 | 0.000947 |
| $y_{26}$-6$H_2O$ | 2429.100 | 2431.307 | 2.207 | 0.000909 |
| $y_{27}$-7$H_2O$ | 2512.147 | 2514.012 | 1.864 | 0.000742 |
| $y_{28}$-7$H_2O$ | 2583.184 | 2585.090 | 1.906 | 0.000738 |
| $y_{30}$-7$H_2O$ | 2757.231 | 2759.044 | 1.813 | 0.000658 |
| $y_{31}$-7$H_2O$ | 2943.310 | 2944.565 | 1.255 | 0.000432 |
| $y_{32}$-8$H_2O$ | 3026.358 | 3028.611 | 2.253 | 0.000749 |
| $y_{33}$-9$H_2O$ | 3109.405 | 3111.700 | 2.295 | 0.000738 |
| $y_{34}$-9$H_2O$ | 3166.427 | 3168.656 | 2.229 | 0.000704 |
| $y_{35}$-9$H_2O$ | 3223.448 | 3225.678 | 2.230 | 0.000692 |
| $y_{36}$-9$H_2O$ | 3294.485 | 3296.715 | 2.230 | 0.000677 |
| $y_{37}$-9$H_2O$ | 3393.554 | 3396.012 | 2.458 | 0.000724 |
| $y_{38}$-9$H_2O$ | 3450.575 | 3453.064 | 2.488 | 0.000721 |
| $y_{39}$-9$H_2O$ | 3565.602 | 3567.832 | 2.230 | 0.000625 |
| $y_{40}$-9$H_2O$ | 3678.686 | 3680.916 | 2.230 | 0.000606 |
| $y_{41}$-9$H_2O$ | 3806.745 | 3808.974 | 2.230 | 0.000586 |
| $y_{42}$-9H2O | 3921.772 | 3924.001 | 2.230 | 0.000569 |
| $y_{43}$-9H2O | 4036.799 | 4039.042 | 2.243 | 0.000556 |

**Table S4. Primers used for cloning of RosM and RosA2α/RosA1β.**

| Primer | Sequence | Source | Identifier |
|---|---|---|---|
| ACYCDuetUP1 | GGATCTCGACGCTCTCCCT | Novagen | Cat#71178-3 |
| DuetDOWN1 | GATTATGCGGCCGTGTACAA | Novagen | Cat#71179-3 |
| DuetUP2 | TTGTACACGGCCGCATAATC | Novagen | Cat#71180-3 |
| T7 Terminator | GCTAGTTATTGCTCAGCGG | Novagen | Cat#69337-3 |
| M1 | GAGGAACGCTTCGCCTCCTA | This paper | N/A |
| M2 | AGGACGGCACCTTCGGCCTG | This paper | N/A |
| M3 | GACCCCGGCGACTTCGCGGA | This paper | N/A |

**Table S5. Locus tag of protein sequences used in this study**

| | |
|---|---|
| Lacticin 3147 | LtnA1, O87236<br>LtnA2, O87237 |
| Staphylococcin C55 | SacA1, BAB78438<br>SacA2, BAB78439 |
| Plantaricin W | PlwA1, AAG02567<br>PlwA2, AAG02566 |
| BhtA | BhtA1, AAZ76603<br>BhtA2, AAZ76602 |
| Lichenicidin | LchA1, P86475<br>LchA2, P86476 |
| Haloduracin | HalA1, Q9KFM5<br>HalA2, Q9KFM6 |
| BhtA | BhtA1, Q3YB75<br>BhtA2, Q3YB76 |
| Enterocin W | EnwA, H3JSS9<br>EnwB, H3JST0 |
